# DGMP: Identifying Cancer Driver Genes by Jointing DGCN and MLP from Multi-Omics Genomic Data

**DOI:** 10.1101/2022.02.16.480791

**Authors:** Shao-Wu Zhang, Jing-Yu Xu, Tong Zhang

## Abstract

Identification of cancer driver genes plays an important role in precision oncology research, which is helpful to understand the cancer initiation and progression. However, most of existing computational methods mainly used the protein-protein interaction networks (PPIs), or treated the directed gene regulatory networks (GRNs) as the undirected gene-gene association networks to identify the cancer driver genes, which will lose the unique structure regulatory information in the directed GRNs, and then affect the outcome of the cancer driver genes identification. Here, based on the multi-omics pan-cancer data (*i.e.*, gene expression, mutation, copy number variation and DNA methylation), we proposed a novel method (called DGMP) to identify cancer driver genes by jointing Directed Graph Convolution Network (DGCN) and Multilayer Perceptron (MLP). DGMP learns the multi-omics features of genes as well as the topological structure features in GRN with DGCN model, and uses MLP to weight more on gene features for mitigating the bias toward the graph topological features in DGCN learning process. The results on three gene regulation networks show that DGMP outperforms other existing state-of-the-art methods. It can not only identify highly mutated cancer driver genes but also the driver genes harboring other kinds of alterations (*e.g.*, differential expression, aberrant DNA methylation) or genes involved in GRNs with other cancer genes. The source code of DGMP can be freely downloaded from https://github.com/NWPU-903PR/DGMP.

## Introduction

Cancer is a heterogeneous disease that is driven by various kinds of genomic and epigenomic alterations, such as single nucleotide variations, DNA methylation, and chromosomal aberrations [1]. Some of these alterations confer growth and positive selection advantages to the mutant cells, leading to intensive proliferation and tumors [2]. That is, the accumulation of diverse genetic mutations causes the cancer progression, and these genetic mutations confer a selective growth advantage to the mutant cells [3]. Thus, identification and comprehensive understanding of cancer driver genes that play causal roles in cancer evolution are crucial for cancer diagnosis and therapy [4].

Several cancer sequencing projects [5–7] have generated a large volume of gene mutation data. Therefore, many computational methods have been proposed to identify the driver genes and disease genes from the cancer genomics data. Generally, these methods can be cataloged into three groups: (1) mutation frequency-based methods, (2) network-based methods, and (3) machine learning-based methods.

Mutation frequency-based methods identify the significantly hypermutated genes as the driver genes compared with a background mutation frequency distribution [8, 9]. For example, MutSigCV [8] calculated the statistical significance of their mutation frequency among all the samples to identify the recurrently mutated genes as the driver genes. However, due to the tumor heterogeneity, it is difficult to build a reliable background mutation model. In addition, these methods cannot be used to detect the low-mutated frequency and non-mutated cancer driver genes, because part of driver genes are mutated at high frequencies (> 20%), while most of cancer genes are mutated at intermediate frequencies (2%–20%) or even lower frequencies [10], and even many genes involved in tumorigenesis are not altered on the DNA sequences, and these genes are dysregulated through various cellular mechanisms [3].

Network-based methods often adopted random walk with restart (RWR) [11, 12], network diffusion [13, 14], subnetwork enrichment analysis [15–17], matrix completion [18] and network structure control [19–22] to predict cancer driver genes and disease genes at the biological network level by incorporating the protein-protein interactions, pathway knowledge, etc. For example, pgWalk [11] constructed a disease-gene network by integrating the multiple genomic and phenomic data, and then simulated the process of a random walker wandering on such a heterogeneous network to prioritize the candidate genes. MAXIF [15] constructed a phenome–interactome network by integrating the given phenotype similarity profile, protein-protein interaction network and associations between diseases and genes, and then maximized the information flow in this phenome–interactome network to uncover the candidate disease genes. Jiang et al [16] constructed a gene semantic similarity network by the biological process domain of the gene ontology, and then used the gene semantic similarity scores in network to infer disease genes. Although these methods have been successfully used for detecting cancer driver genes and disease genes, they are still limited to the unreliable and incomplete interactions in biological network [23]. Developing an integrative framework by incorporating cancer multi-omics data (*e.g.*, somatic mutations, structural variations, gene expression and methylation) and adopting the hybrid approaches would improve the prediction of cancer driver genes for the network-based methods.

Machine learning-based methods [24–28] usually train the classifier, *e.g.*, random forest and support vector machine (SVM), by extracting the diverse features from different types of cancer data to predict new cancer driver genes. For example, deepDriver [26] predicted cancer driver genes with a convolutional neural network (CNN) model that was trained with the gene mutation features and their neighbors in the similarity networks. Integrating the functional impact of mutations and the similarity of gene expression patterns with CNN model can improve the prediction accuracy of driver genes. NRFD [28] constructed a cancer gene interaction network by integrating various kinds of cancer-related information sources to obtain the feature vector of each gene, and then used the random forest to predict the cancer driver genes.

Most existing machine learning-based methods just extract the network-based features by using the network analysis. They cannot effectively combine the network topology features and the gene multi-omics features. That is, very few methods can combine both multidimensional gene features with the graph representation features of gene-gene interaction networks. For example, EMOGI [29] adopted the graph convolution network (GCN) model to combine the multidimensional multi-omics gene features with the topological features of the protein-protein interaction (PPI) network to identify cancer genes. Although EMOGI [29] successfully identified not only highly mutated cancer genes but also other non-mutated cancer-dependency genes, it only used the association information between genes in undirected PPI network, and does not make full use of the regulation information between genes in gene regulatory network(GRN). In addition, the spectral-based GCN can only be applied to the undirected network. While GRN is a directed network that provides the specific causal links (*e.g.*, one gene activates or inhibits other genes) between genes, which helps to understand the molecular mechanism of gene regulation in cancers and the molecular basis of cancer subtypes [30, 31]. Thus, we proposed a novel deep learning-based method (called DGMP) to identify the cancer driver genes by integrating the multidimensional multi-omics gene features as well as the topological structure features of the GRN through the directed graph convolution network (DGCN) model. Compared to the graph convolution network (GCN), DGCN [32] uses the first- and second-order proximity to extend the spectral-based graph convolution to directed graphs for retaining the connection properties in the directed graph, and also expanding the receptive field in convolution operation without stacking more convolution layers.

Generally, the role of graph in GCN is to guide the weights training by averaging the features of a node with its graph neighbors [33]. When the relationships represented through the graph are consistent with the information of nodes (*i.e.*, the features of neighbor nodes in graph are expected to be more similar than that of other nodes), GCN can improve the performance of nodes classification [33, 34]. If feature similarities of neighborhood nodes in graph are not congruent, this graph-based averaging is not beneficial in the training process. It will result in the performance of GCN lower than that of these methods (*e.g.*, multilayer perceptron) that rely exclusively on the node feature information [33, 34]. Considering that the features of some genes that are graph neighbors in GRN may not be more similar than other genes, we also introduced the multilayer perceptron (MLP) classifier into DGMP model for further improving the performance of cancer driver gene prediction. DGMP uses DGCN model to learn the multi-omics features of genes, as well as the topological structure features in GRN, and adopts MLP to weight more on gene features for mitigating the bias toward the graph topological features in DGCN learning process. DGMP aims to not only identify the highly mutated cancer driver genes, but also predict the driver genes that harbor other kinds of alterations (*e.g.*, differential expression, aberrant DNA methylation), or identify the driver genes involved in GRN with other cancer genes. Overall, our main contributions include two points: (1) We use DGCN model to perform the graph convolution operation on directed gene regulatory network (GRN) for capturing the directed information (*i.e.*, regulation information), and expand the receptive field to second-order neighbor of a gene for aggregating more its neighbor and the topological feature information. (2) We introduce MLP into DGMP framework to weight more on gene features for mitigating the bias toward the topological features in DGCN learning process.

The results on three networks of DawnNet, KEGG and RegNetwork show that our DGMP outperforms other existing state-of-the-art methods in terms of the area under the ROC-curve (AUC) and the area under the precision-recall curve (AUPR), demonstrating that our DGMP can effectively identify the cancer driver genes.

## Method

### DGMP model

DGMP is based on directed graph convolution network (DGCN) and multilayer perceptron (MLP), and trained in a semi-supervised manner to discriminate the cancer driver genes from the non-cancer driver genes. The inputs of DGMP are gene single nucleotide variants (SNVs), gene copy number aberrations (CNVs), gene expression information, DNA methylation in gene promoter regions, and the gene regulatory network (GRN) in which some genes have labels, while most have no labels. The positive labels corresponding to the annotated cancer driver genes, and negative labels corresponding to the non-caner driver genes in the partially labelled GRN. The output of DGMP is a fully labelled graph, in which each gene is assigned a probability to be a cancer driver gene. As shown in Figure 1, DGMP mainly consists of three modules. The first module (Figure 1A) is that DGCN [32] is used to learn the embedding vectors of genes from GRN and genomic data (*i.e.*, SNVs, CNVs, DNA methylation and gene expression) by utilizing the first-order proximity (*A*_*F*_), the second-order in-degree proximity (*A*_*in*_) and the second-order out-degree proximity (*A*_*out*_). The first- and second-order proximity can expand the convolution operation receptive field, extract and leverage the directed graph information. The second module (Figure 1B) is the multilayer perceptron (MLP) which is used to obtain the embedding vectors of genes only from the genomic data of SNVs, CNVs, gene expression and DNA methylation. The third module (Figure 1C) is a fully connected neural network which is built to predict the cancer driver genes by concatenating the output embedding vectors from DGCN and MLP.

**Figure 1.**
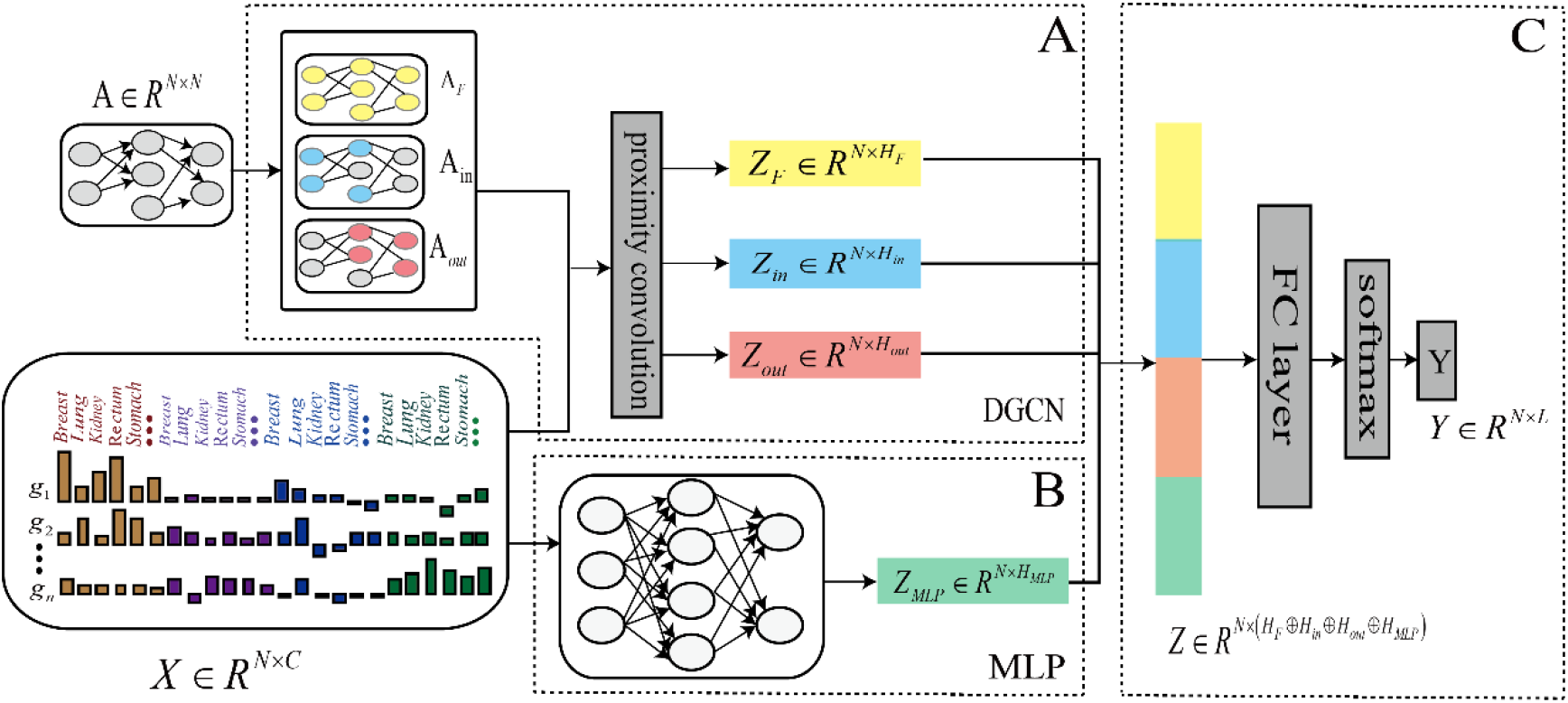
Schematic of DGMP model framework. **A.** DGCN module. GRN with partial labelled genes and the genomic information (i.e., SNVs, CNVs, DNA methylation and gene expression) are inputted into the DGCN module. According to the definition of first-order and second-order proximity, we can obtain three proximity networks (i.e., three undirected graphs) of the first-order proximity network (A_F_), second-order in-degree proximity network (A_in_) and second-order out-degree proximity network (A_out_) from GRN (i.e., directed graph), and then implement graph convolution operation on these three proximity networks to achieve graph convolution of directed graph for generating three embedding vectors (Z_F_, Z_in_ and Z_out_) of genes. **B.** MLP module. The genomic data of SNVs, CNVs, gene expression and DNA methylation are inputted into the multilayer perceptron (MLP) to obtain the embedding vectors (ZMLP) of genes. **C.** Fully connected neural network module. The first-order proximity convolution output Z_F_, second-order in-degree proximity convolution outputs Z_in_, and second-order out-degree proximity convolution outputs Z_out_ of DGCN, the output ZMLP of MLP are concatenated in series to form an embedding matrix Z that is fed into a fully connected neural network to predict the cancer driver genes. *g*_*n*_ represents the *n*-th gene; *N* is the total number of genes; *C* represents the dimension of gene features; *L* represents the dimension of gene labels; *H*_*F*_, *H*_*in*_, *H*_*out*_ and *H*_*MLP*_ represent the dimension of gene embedding feature vectors.

In the process of training our DGMP model, the semi-supervised training manner is implemented in DGCN module. By inputting both the structural features of partially labelled GRN and the multi-omics features of genes, DGCN encodes GRN structure by directly using a neural network model to obtain the embedding feature vectors of all genes in GRN (in which some genes have labels, while most have no labels), and then trains on a supervised target for all genes with labels. That is, all the labeled and unlabeled genes participate in the generation of graph embedding vectors, and only the labeled genes are used to evaluate the loss of DGMP.

### Directed Graph Convolution Network (DGCN)

In order to use the graph convolutional network (GCN) model which can effectively learn the underlying pairwise relationship among vertices in directed graph, Tong et al. [32] used the first-and second-order proximity to extend the spectral-based graph convolution to the directed graphs, and then developed a directed graph convolution network (DGCN) model to learn the embedding vectors of nodes in directed graphs.

For a directed gene interaction network *G*, it can be considered a multi-layer graph convolutional network by using first-and second-order proximity of the network *G* [32].

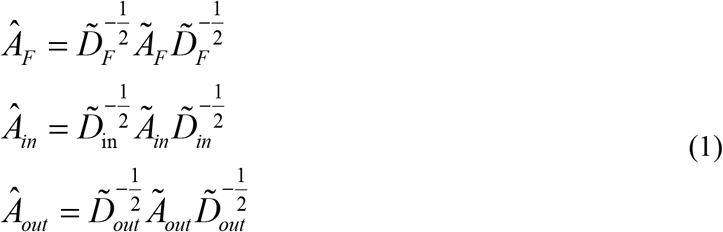

and

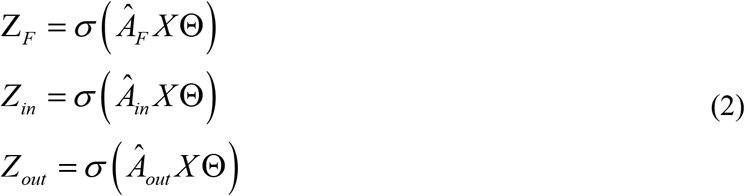

where σ(•) is an activation function; *Ã*_*F*_ is the first-order proximity matrix with self-loop derived from the gene interaction network matrix *A* ∈ *R*^*N* × *N*^ with partial labelled genes (*i.e.*, cancer driver genes, non-cancer driver genes and unlabeled genes); *Ã*_*in*_ is the second-order in-degree proximity with self-loop derived from the gene interaction network matrix *A*, *Ã*_*out*_ is the second-order out-degree proximity with self-loop derived from the gene interaction network matrix *A*; *X* is the feature matrix of genes; Θ is a shared trainable weight matrix; *Z*_*F*_ is the first-order proximity convolution output of DGCN; *Z*_*in*_ and *Z*_*out*_ are the second-order in- and out-degree proximity convolution outputs of DGCN. *Z*_*F*_, *Z*_*in*_ and *Z*_*out*_ can not only obtain the first and second-order neighbor feature information in network *G*, but also *Z*_*in*_ and *Z*_*out*_ retain the directed structure information in network *G*.

The first-order proximity entry *A*_*F*_ (*i*, *j*), the second-order in-degree proximity entry *A*_*in*_ (*i*, *j*), and the second-order out-degree proximity entry *A*_*out*_ (*i*, *j*) between gene *v*_*i*_ and *v*_*j*_ in matrix *A* are defined as follows:

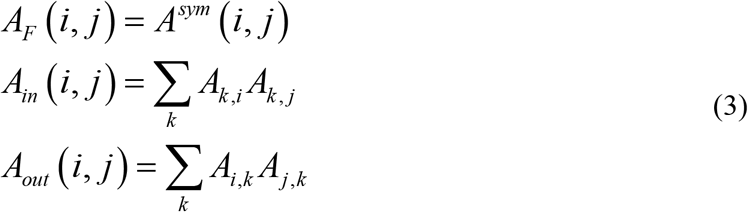

where *A*^*sym*^ is the symmetric matrix of matrix *A*. If there is no link from *v*_*i*_ to *v*_*j*_ or *v*_*j*_ to *v*_*i*_, then *A*_*F*_ (*i*, *j*) = 0. *A*_*in*_(*i*, *j*) is an entry in the in-degree matrix of matrix *A*. *A*_*out*_ (*i*, *j*) is an entry in the out-degree matrix of matrix *A*.

Since *A*_*in*_ (*i*, *j*) is the sum of the input edges both of gene *v*_*i*_ and *v*_*j*_, i.e., Σ_*k*_*A*{*i* ← *k* → *j*} which reflects the similarity of the in-degree between gene *v*_*i*_ and *v*_*j*_. The greater the value of *A*_*in*_(*i*, *j*), the higher the similarity of the second-order in-degree. Similarly, *A*_*out*_ (*i*, *j*) measures the second-order out-degree proximity by accumulating the links from both gene *v*_*i*_ and *v*_*j*_, i.e., Σ_*k*_*A*{*i* ← *k* → *j*}. If there are no shared genes linked from gene *v*_*i*_ to *v*_*j*_, their second-order proximity is set to zero.

### Multilayer Perceptron (MLP)

Multilayer Perceptron (MLP) is a forward-structured artificial neural network, which maps a set of input vectors to a set of output embedding vectors. The feature matrix *X* of genes is inputted into the MLP to get the embedding matrix *Z*_*MLP*_.

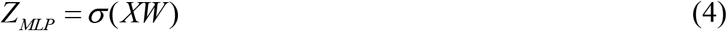

where *X* is the feature matrix of genes, *W* is a shared trainable weight matrix, and σ(.) is an activation function.

### Fully connected neural network

The first-order proximity convolution output *Z*_*F*_, the second-order in-degree proximity convolution output *Z*_*in*_, the second-order out-degree output *Z*_*out*_ from DGCN, and the output matrix *Z*_*MLP*_ from MLP are concatenated to form a embedding matrix *Z*, which is fed into a fully connected neural network to obtain the probability of a gene as a cancer driver gene.

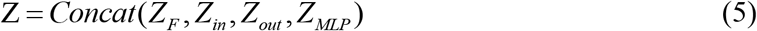

In summary, DGMP model can be written as follows.

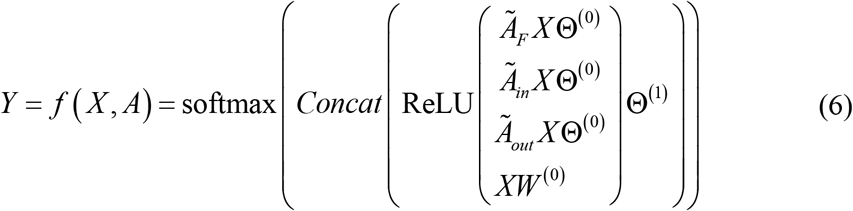

Three different proximity convolutions on GRN share a same filter weight matrix *Θ*^(0)^ ∈ ℝ^*C*×*H*^. It transforms the input dimension *C* to the embedding size *H*. The weight matrix *W*^(0)^ ∈ ℝ^*C*×*H*^ in MLP also transforms the input dimension *C* to the embedding size *H*. The outputs of three different proximity convolutions on GRN and the output of MLP are concatenated to feed into a fully connected network layer, which is used to convert the feature dimension from 4*H* to *F*. Θ^(1)^ ∈ ℝ^4*H*×*F*^ is an embedding-to-output weight matrix. The activation function is defined as *softmax*(*x*_*i*_) = *exp*(*x*_*i*_)/Σ_*i*_ = *exp*(*x*_*i*_).All labelled samples are used to calculate the cross-entropy error in this semi-supervised cancer driver gene identification task.

We utilized ADAM optimizer for training 500 epochs, and set the dropout rate to 0.6, learning rate 0.001, weight decay 0.01. In our DGMP model, the dimensions of all the four embedding vectors (i.e., Z_F_, Z_in_, Z_out_ and Z_MLP_) were set to 4.

## Results

### Datasets

The genomics data of gene expression, gene mutation, copy number and DNA methylation used in EMOGI work are took to assess the performance of our DGMP model. These genomics data are collected from 29,446 samples in the TCGA database, covering 16 different cancer types. By performing the same data preprocessing pipeline as EMOGI, we can obtain the pan-cancer gene feature matrix *X* (*X* ∈ *R*^*N* × 64^) in which each gene is represented with a 16×4 dimensional vector, here *N* is the gene number, 16 is the number of cancer types, and 4 refers to the values of four omics types (i.e., single nucleotide variants, copy number aberrations, DNA methylation and gene expression) that are computed for each cancer type.

The gene regulatory networks of DawnNet, KEGG and RegNetwork are selected to implement our DGMP model and other competitive methods. For the GRN of DawnNet built in DawnRank [35], we merge all the redundant genes into single genes, and combine their corresponding edges to generate a directed gene-gene association network (namely DawnNet) that contains 9,677 genes, 176,826 directed edges and 10,150 undirected edges. For the KEGG network, we integrated the pathways in KEGG database [36]with Pathview tool [37] to obtain the KEGG pathway network. KEGG network contains 4,798 genes and 61,520 directed edges. For the RegNetwork, we extracted the regulatory interactions of TF-TF and TF-gene from RegNetwork data repository [38] to generate a directed network, named as RegNetwork. RegNetwork contains 20,300 TFs/genes, 148,387 regulation interaction edges. RegNetwork data repository [38] was established by integrating the documented regulatory interactions among transcription factors (TFs), microRNAs (miRNAs) and target genes from 25 selected databases, including five-type transcriptional and posttranscriptional regulatory relationships (*i.e.*, TF-TF, TF-gene, TF-miRNA, miRNA-TF, miRNA-gene) for human and mouse. The data of DawnNet, KEGG and RegNetwork can be downloaded from Tables S1-S3 (in supplementary B).

The known cancer driver genes are obtained from the expert-curated list in NCG [6] and IntOGen[39] to form a positive set *S*^+^. The non-cancer driver genes can be selected from those most likely to be unassociated with cancers. We use the following criteria to recursively remove the genes from the set of all genes to get the non-cancer driver genes, thus forming a negative set *S*^−^. (1) Removing the genes that are part of the known cancer driver genes in NCG; (2) Removing the genes that present in the OMIM disease database [40]; (3) Removing the genes that associate with cancer pathways in the KEGG database [36]; (4) Removing the genes whose expression is correlated to the expression of cancer driver genes [41]. Generally, the number of non-cancer driver genes is far more than that of the known cancer driver genes. In order to avoid the bias of the prediction model towards the negative samples in the training process, we randomly sample the non-cancer driver genes from the negative set *S*^−^, whose number are same as that of the known cancer driver genes. In addition, only the positive and negative samples that are included in the directed networks of DawnNet, KEGG and RegNetwork are used for training. That is, we used 693 positive samples and 693 negative samples to train the prediction models for DawnNet network, and 406 positive samples and 406 negative samples for KEGG network, and 826 positive samples and 826 negative samples for RegNetwork network.

### Performance comparison of DGMP with other methods

In this work, we take the AUC and AUPR metrics to evaluate the prediction power of different methods. AUC value is defined as the area under the receiver operating characteristic (ROC) curve, which plots the false positive rate (FPR) against the true positive rate (TRP) at different thresholds. AUPR value is defined as the area under the precision-recall (PR) curve, which plots the ratio of true positives among all positive predictions for each given recall rate. AUPR is a more significant quality metric than AUC for identifying the cancer driver genes, because it punishes much more the existence of false positive cancer driver genes among the best ranked prediction scores [42].

To evaluate the performance of our DGMP for identifying the pan-cancer driver genes, we first compared our DGMP with the machine learning-based methods of NRFD [28], EMOGI [29] and DeepWalk [43] +SVM on DawnNet network in 5-fold cross-validation (5CV) test, and then compared it with the network-based methods of PageRank [44] and HotNet2 [13], and the mutation frequency-based method of MutSigCV [8]. For 5CV test [45], all the labelled genes (*i.e.*, known cancer driver genes and the non-cancer driver genes selected according to some criteria) are randomly partitioned into 5 non-overlapping subsets of roughly equal size. One of these subsets is singled out in turn as the test set and other four subsets are used as the training sets. This process is repeated for 5 iterations until all the labelled genes are tested in turn. In addition, we also performed these methods on KEGG and RegNetwork networks. The results of our DGMP and other six methods on three directed networks of DawnNet, KEGG and RegNetwork are shown in Table 1. The ROC curves and precision-recall curves of these seven methods on DawnNet, KEGG and RegNetwork networks are shown on Figures S1-S6 (in supplementary A), respectively.

**Table 1.**
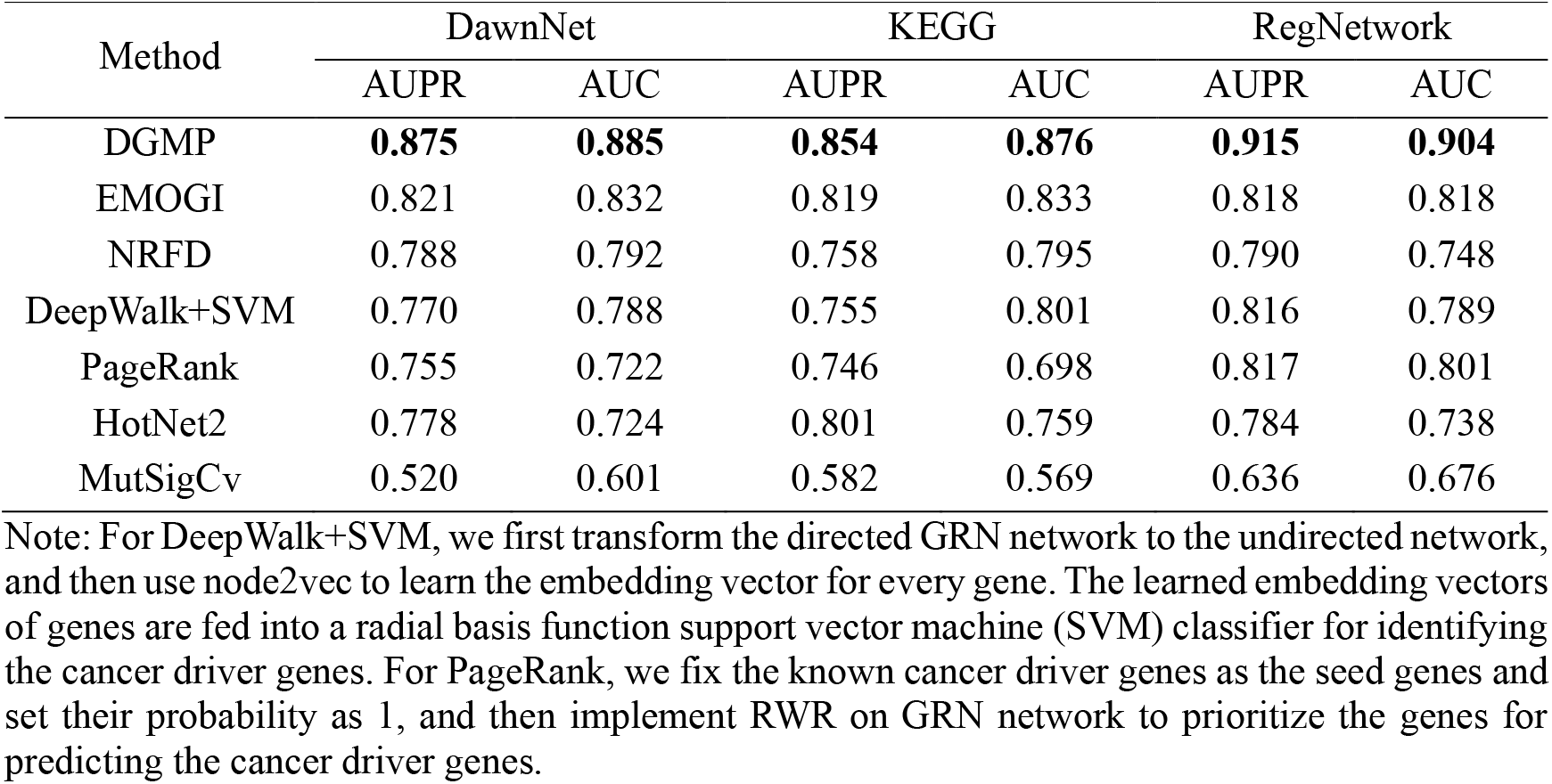
AUPR and AUC of DGMP, EMOGI, NRFD, HotNet2, MutSigCV, PageRank and DeepWalk+SVM on three networks of DawnNet, KEGG pathway and RegNetwork in 5CV test

From Table 1 and Figures S1-S6, we can see that the performance of our DGMP outperforms all of the other six state-of-the-art methods of EMOGI, NRFD, DeepWalk+SVM, HotNet2, PageRank and MutSigCV for identifying the cancer diver genes on three directed networks of DawnNet, KEGG and RegNetwork. The AUPR and AUC values of DGMP on DawnNet network are 0.875 and 0.889, which are 0.054~0.355, 0.053~0.284 higher than that of other six methods, respectively; AUPR and AUC values of DGMP on KEGG pathway network are 0.854 and 0.876, which are 0.035~0.272, 0.043~0.307 higher than that of other six methods, respectively; AUPR and AUC values of DGMP on RegNetwork network are 0.915 and 0.904, which are 0.097~0.279, 0.086~0.228 higher than that of other six methods, respectively. These results demonstrate that our DGMP method has superior performance in identifying the cancer driver genes.

To further assess the performance of DGMP, we designed other two scenarios of using the unbalanced positive and negative training samples to train the prediction models on DawnNet and STRING PPI [46] networks. That is, we used 693 cancer driver genes and 1,763 non-cancer driver genes to train the prediction models on DawnNet network, and utilized 734 cancer driver genes and 1,152 non-cancer driver genes to train the prediction models on STRING PPI network (listed in Table S4 in supplementary B). The experimental results of two scenarios in 5CV test are shown in Table S5 (in supplementary A), from which we can see that DGMP achieves the best performance on two networks in terms of AUC and AUPR values compared with all other methods, demonstrating the effectiveness of DGMP for identifying the cancer driver genes.

In order to evaluate the generalization performance of our DGMP, we built an independent test set of the cancer driver genes from CancerMine database [47]. CancerMine is a literature-mined resource of cancer-related genes, which collects the genes of drivers, oncogenes and tumor suppressors in different types of cancer. The genes in independent test set (listed in Table S6 in supplementary B) do not overlap with those in the training set. To calculate the AUPR and AUC values, we counted hits in the independent test set as true positive samples, and all other predicted genes not contained in the independent test set as the false positive samples. The results of our DGMP and other comparison methods in independent test are shown in Table 2, from which we can see that DGMP achieves the best performance on DawnNet and RegNetwork networks compared with all other methods. Although AUPR and AUC values of DGMP on KEGG network are slightly lower than those of DeepWalk+SVM method, the TPR (True Positive Rate) of DGMP is higher than that of deepwalk+SVM method. For independent test that only know the driver gene labels, TPR is more objective than AUPR and AUC to measure the generalization performance of prediction models, because we counted all other unlabeled driver genes as the non-cancer driver genes in the process of calculating AUPR and AUC, while all these unlabeled (or unannotated) genes are not really non-cancer driver genes. In addition, AUPR and AUC values of DeepWalk+SVM on KEGG network are slightly higher than those of our DGMP, the reason may be that DeepWalk+SVM obtains better performance on smaller network, while our DGMP achieves superior performance on larger networks. These results in independent test further demonstrate the power of DGMP to identify the cancer driver genes.

**Table 2.** Results of DGMP, EMOGI, NRFD, HotNet2, MutSigCV, PageRank and DeepWalk+SVM on DawnNet, KEGG and RegNetwork networks in independent test

| Method | DawnNet |  |  | KEGG |  |  | RegNetwork |  |  |
| --- | --- | --- | --- | --- | --- | --- | --- | --- | --- |
|  | AUPR | AUC | TPR | AUPR | AUC | TPR | AUPR | AUC | TPR |
| DGMP | **0.136** | **0.706** | **0.769** | 0.107 | 0.635 | **0.801** | **0.115** | **0.757** | **0.864** |
| EMOGI | 0.107 | 0.659 | 0.747 | 0.100 | 0.602 | 0.767 | 0.103 | 0.684 | 0.847 |
| NRFD | 0.122 | 0.659 | 0.706 | 0.092 | 0.594 | 0.781 | 0.092 | 0.698 | 0.843 |
| DeepWalk+SVM | 0.124 | 0.696 | 0.671 | **0.119** | **0.659** | 0.790 | 0.098 | 0.697 | 0.662 |
| PageRank | 0.101 | 0.589 | - | 0.089 | 0.545 |  | 0.069 | 0.648 |  |
| HotNet2 | 0.079 | 0.512 | - | 0.101 | 0.538 |  | 0.055 | 0.555 |  |
| MutSigCv | 0.053 | 0.491 | - | 0.073 | 0.483 |  | 0.031 | 0.447 |  |

Note: Since the outputs of PageRank, HotNet2 and MutSigCV methods are the rank orders of genes, not gene labels, we cannot calculate their TPRs. $TPR (True\ Positive\ Rate) = TP/(TP+FN)$ .

### Ablation experiments of diverse architecture components in DGMP

To evaluate the contributions of diverse architecture components in our DGMP, we conducted the ablation experiments on DawnNet network in 5CV test. The ablation experimental results of DGMP are shown in Table 3. In Table 3, DGMP-_direction_ denotes that we neglect the directionality of regulation edges in DawnNet directed network, and adopt DGMP to identify the cancer driver genes. MLP denotes that we remove the DGCN module from DGMP model architecture to identify the cancer driver genes; DGCN denotes that we remove the MLP module from DGMP model architecture to identify the cancer driver genes; DGCN-_direction_ denotes that we neglect the directionality of regulation edges in DawnNet directed network, and adopt DGCN to identify the cancer driver genes. DGCN-_X_ denotes that we just use the topological information of GRN in DGCN module without considering the genomic information (*i.e.*, SNVs, CNVs, DNA methylation and gene expression) of genes, and also remove the MLP module from DGMP model architecture to identify the cancer driver genes; DGCN-_X_MLP denotes that we combine DGCN_-X_ with MLP and the full connected neural network to identify the cancer driver genes.

**Table 3.** The ablation experimental results of Driver_DGCNMP on DawnNet network in 5CV test.

|  | AUPR | AUC |
| --- | --- | --- |
| DGMP | **0.875 ±0.020** | **0.885±0.019** |
| DGMP<sub>-direction</sub> | 0.871±0.0017 | 0.881±0.018 |
| MLP | 0.801 ±0.017 | 0.839±0.018 |
| DGCN | 0.828±0.018 | 0.856±0.022 |
| DGCN<sub>-direction</sub> | 0.825±0.019 | 0.838±0.018 |
| DGCN<sub>-x</sub> | 0.758±0.013 | 0.743±0.016 |
| DGCN<sub>-x</sub>MLP | 0.841 ±0.021 | 0.857±0.016 |

As shown in Table 3, we can see that AUPR of DGMP that consists of DGCN and MLP is 0.875, which is 0.047, 0.074 higher than that of DGCN and MLP respectively, indicating that jointing DGCN and MLP can improve the performance of identifying the cancer driver genes. AUPR of DGCN-XMLP is 0.083 higher than that of DGCN_-X_, indicating that MLP does mitigate the bias toward the graph topological features in DGCN learning process, further enhancing the prediction performance of DGMP. AUPR of DGCN is 0.07 higher than that of DGCN_-X_, indicating that feeding the multi-omics information of genes into DGCN can improve the prediction performance of DGMP. AUPR of DGMP-_direction_ is 0.004 lower than that of DGMP, and DGCN-_direction_ is 0.003, 0.05 lower than that of DGCN and DGMP respectively, demonstrating that considering the regulation information (*i.e.*, directionality of regulation edges) can improve the prediction performance of DGMP. The results in Table 3 show that all the proposed components for building DGMP are valid and contribute to the final performance of DGMP.

Previous studies [33] and [34] show that if the features of neighbors of a center node are not similar, further graph convolution operation will result in lower performance for GCN. Considering that the neighbors’ features of some center genes in GRN may not be more similar than other genes, performing graph convolution operation on these neighbors will reduce the performance of DGCN, we introduce MLP to offset the performance degradation of DGCN. In order to further show the effectiveness of MLP mitigating the bias toward the graph topological features in DGCN learning process, the neighborhood discrete entropy *Scoreetp*_*etp*_ (*u*) [34] (its definition and formulation are given in supplementary A) was used to measure the diversity of a gene’s neighborhood. We picked top 50 predicted cancer driver genes and top 50 predicted non-cancer driver genes with the highest discrete entropy, and then employed t-SNE to visualize the distribution (Figure 2) of these genes by extracting their embedding vectors (*i.e.*, *Z*_*F*_, *Z*_*in*_, *Z*_*out*_ and *Z*_*MLP*_). From Figure 2, we can see that after concatenating these embedding vectors (*i.e.*, *Z*_*F*_, *Z*_*in*_, *Z*_*out*_) generated by DGCN with the embedding vectors (i.e., *Z*_*MLP*_) generated by MLP, the within-class variance of cancer/non-cancer driver genes is smaller than that of DGCN, and the between-class distances of cancer/non-cancer driver genes are larger than that of DGCN in embedding space. These results demonstrate that graph convolution operation is not so good for distinguishing these genes when the features of their neighbors are dissimilar, while MLP can offset the performance degradation of DGCN, that is, MLP can mitigate the bias toward the graph topological features in DGCN learning process.

**Figure 2.**
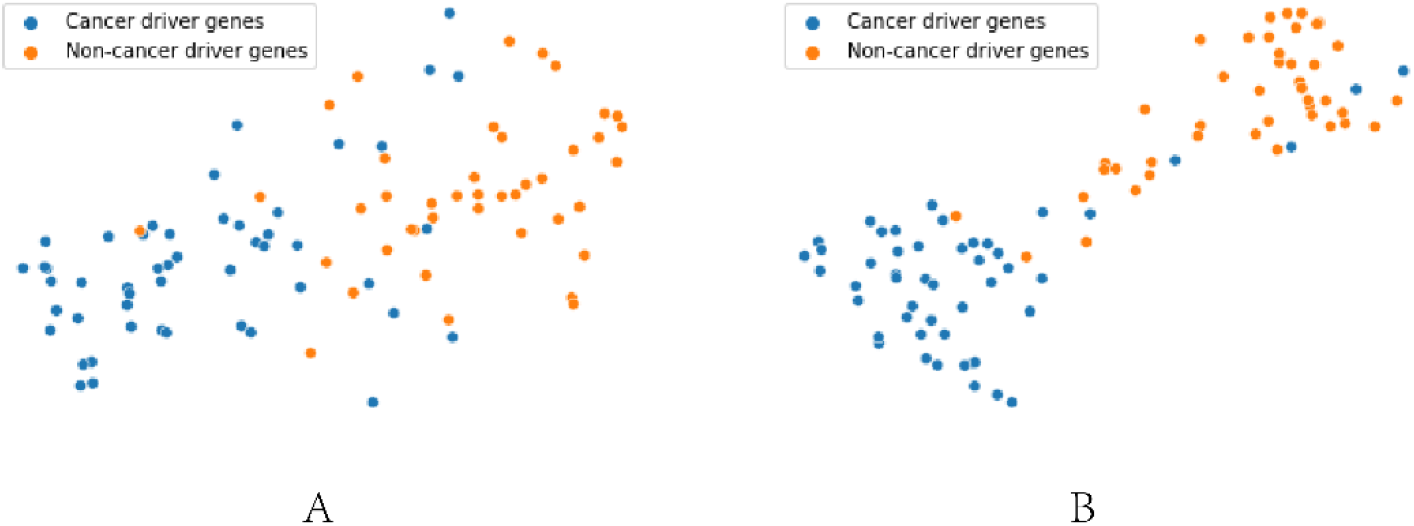
t-SNE visualization of top 50 predicted cancer/non-cancer driver genes with high neighbor discrete entropy. **A.** Visualization of 50 cancer/non-cancer driver genes by concatenating *Z*_*F*_, *Z*_*in*_ and *Z*_*out*_ from DGCN. **B.** Visualization of 50 cancer/non-cancer driver genes by concatenating *Z*_*F*_, *Z*_*in*_, *Z*_*out*_ from DGCN and *Z*_*F*_, *Z*_*in*_ and *Z*_*MLP*_ from MLP.

### De novo cancer driver gene analysis

To analyze the de novo cancer driver genes, we selected the genes that are not annotated as driver genes in NCG database [6] from top100 cancer driver genes (see Table S7 in supplementary B) predicted by DGMP on DawnNet, and considered these genes as the newly predicted cancer driver genes (namely NPCGs). As a result, we obtained 52 NPCGs (listed in Table S8 in supplementary B), of which 41 NPCGs are labelled as the cancer driver genes, or oncogene/tumor suppressor genes in CancerMine database [47]. For example, transcriptional activator SP3 as a driver gene competes with the tumor suppressor AP-2 for binding the VEGF promoter in prostate cancer, thereby repressing AP-2 expression [48]; TFF2 expression inhibits the gastric cancer cell growth and invasion in vitro via interactions with the transcription factor Sp3, and Sp3 knockdown in gastric cancer cells antagonizes TFF2 antitumor activity [49]. NFKB1 is a tumor suppressor in cervical cancer by inhibiting cell proliferation, colony formation and migration, and its mutation will affect the radiotherapy sensitivity in cervical cancer [50]. GBP1 is downregulated and acts as a tumor suppressor in colorectal cancer cells [51]. These evidences show that our DGMP can effectively predict the new and candidate cancer driver genes. We also designed the following experiments to demonstrate the effectiveness of our DGMP in predicting de novel cancer driver genes.

Firstly, we calculated the interaction percentages of known cancer driver genes (KCGs) from NCG database [6] with NPCGs, candidate cancer driver genes (CCGs) [6] and KCGs, as well as the interaction percentage of KCGs with other genes that neither belong to NPCGs nor KCGs and CCGs. The statistical results are shown in Figure 3A. As shown in Figure 3A, we can find that genes in NPCGs generally have more interactions with KCGs than other genes, indicating that the NPCGs predicted by DGMP are closely related to initiation and progression of cancers. We also found that SP1 and SHC1 have the largest number of interactions with KCGs. Among them, SP1 strongly associated with Ser345-phosphorylated PR-B receptors to regulate growth-promoting (EGFR) target genes and PR cell cycle (p21) for breast cancer cell proliferation [52]; SHC1 may be an important route of DEPDC1B regulating the development of bladder cancer. In DEPDC1B-overexpressed cancer cells, the knockdown of SHC1 could abolish the promotion effects caused by DEPDC1B [53]. In addition, we also compared the degrees of genes in NPCGs, KCGs and CCGs with the degrees of other genes that neither belong to NPCGs nor KCGs and CCGs. As shown in Figure 3B, we can see that the degrees of genes in NPCGs, KCGs and CCGs are significantly larger than that in other gene set, indicating that the greater the network degree of a gene, the more likely it is to be as a cancer driver gene.

**Figure 3.**
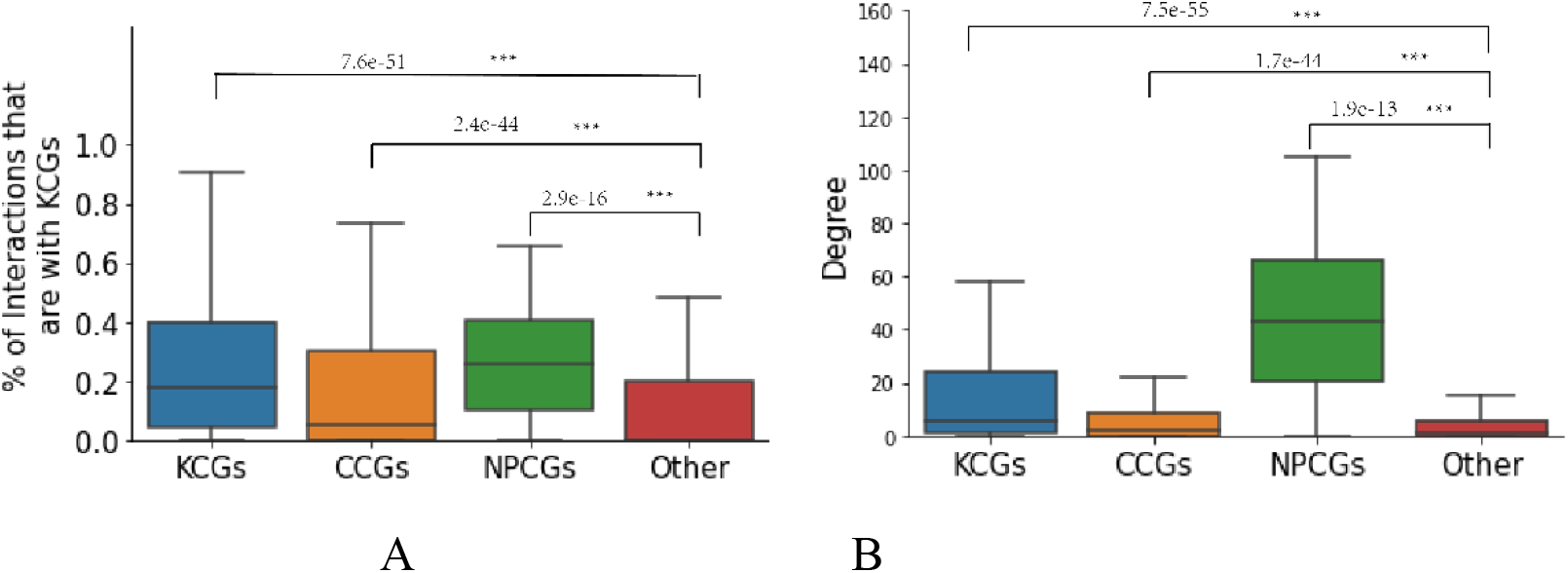
Statistical results of other genes, NPCGs, KCGs and CCGs interacting with KCGs. **A.** Percentages of other genes, NPCGs, KCGs and CCGs interacted with KCGs. **B.** Degree of other genes, NPCGs, KCGs and CCGs interacted with KCGs. Mann-Whitney U test was carried out to test the difference significance of each comparison.

Secondly, we analyzed the multi-omics gene features of NPCGs, KCGs, CCGs and other genes by calculating the average frequency of SNVs, average CNVs rate, average DNA methylation changes and average gene differential expression changes across 16 cancer types. The mutation types contains both truncating and gain-of-function mutations, and all SNVs have the potential to affect cell growth.

As shown in Figure 4, we found that there are significant differences in SNV mutations (p-value=2.1e-04), CNVs (p-value=0.04), gene differential expression (p-value=4.3e-03) between other genes and NPCGs, while there is no significant difference (p-value=0.2) between other genes and NPCGs for DNA methylation. These results indicate that the omics gene features of SNVs, CNVs and gene differential expression are the important factors in distinguishing NPCGs from other genes. The NPCGs identified by DGMP are more frequently mutated across samples than other genes, indicating that DGMP can effectively identify the cancer driver genes from highly mutated genes. For example, EGF and ROCK1 are the high mutation rate genes across different cancer types in NPCGs, which are demonstrated to be correlated with cancers [47, 48]. EGF enhances the phosphorylation and acetylation of histone H3 to promote the DKK1 transcription in hepatocellular carcinoma[54]; ROCK1 is over-expressed in human hepatocellular carcinoma (HCC) cell lines and tissues, and knockdown of ROCK1 or ROCK2 can inhibit the HCC cell growth [55]. In addition, NPCGs also contains highly DNA methylation genes and highly mRNA differential expressed genes, which rarely mutate across cancers. For example, ITGB3 gene is highly DNA methylation but with relatively lower SNVs frequency, whose high expression is correlated with overall survival and worse progression-free survival of multiple myeloma patients [56]; TFAP2A gene is highly overexpressed compared to normal tissue in multiple cancer types, which modulates ferroptosis in gallbladder carcinoma cells through the Nrf2 signaling axis [57]. These results show that our DGMP can not only identify the driver genes involved in GRN with other known cancer genes, but also the highly mutated cancer driver genes or driver genes harboring other kinds of alterations (*e.g*., differential expression, aberrant DNA methylation).

**Figure 4.**
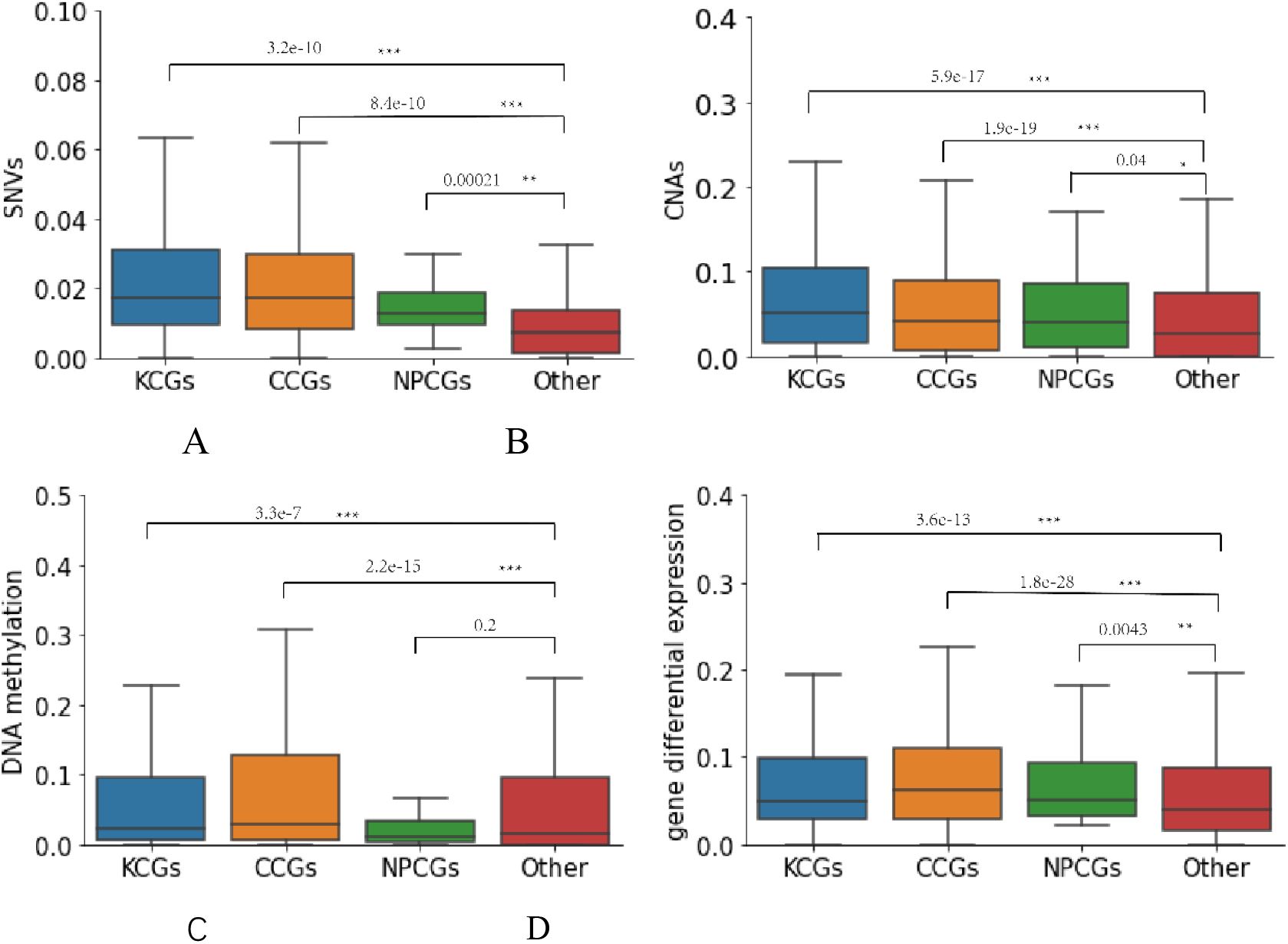
Averaged SNVs, CNVs, DNA methylation, gene differential expression across 16 cancer types for other genes, NPCGs, KCGs and CCGs. **A.** Average frequency of SNVs. **B.** Average CNVs rate. **C.** Average DNA methylation change. **D.** Average gene differential expression change. Mann-Whitney U test was carried out to test the difference significance of each comparison

## Conclusions

In this work, we presented a novel method of DGMP to identify cancer driver genes by integrating multi-omics genomic data and jointing the directed graph convolution network (DGCN) and multi-layer perceptron (MLP). DGMP uses DGCN to learn the multi-omics features of genes as well as the topological structure features of gene regulation network (GRN), and employs MLP to learn the gene features from multi-omics genomic data. Three embedding feature vectors from DGCN and one embedding feature vector from MLP are concatenated to feed into a fully connected neural network to output the probability that one gene is a cancer driver gene. The results on three networks of DawnNet, KEGG and RegNetwork show that DGMP is superior to other existing state-of-the-art methods in identifying the cancer driver genes. The analysis results of top100 predicted cancer driver genes demonstrate that DGMP not only identifies more known caner driver genes, but also can effectively predict the highly mutated cancer driver genes, and the driver genes harboring other kinds of alterations (*e.g.*, differential expression, aberrant DNA methylation), or genes involved in GRN with other cancer genes. The t-SNE visualization distribution of 50 cancer/non-cancer driver genes with high neighbor discrete entropy show that concatenating the embedding feature vectors of DGCN and MLP can improve the aggregation of cancer/non-cancer driver genes, that is, MLP indeed mitigates the bias toward the graph topological features in DGCN learning process. In addition, DGMP can also be used to successfully identify driver genes of the specific caners, such as breast cancer and thyroid cancer (see Figure S7 in supplementary B).

There are two potential reasons which are responsible for the remarkable performance of DGMP. The first aspect is that DGMP makes full use of the regulation information among genes by adopting the DGCN model. The second aspect is that DGMP introduces MLP to weight more on gene features for mitigating the bias toward the graph topological structure features in DGCN learning process, which offsets the performance degradation of DGCN caused by the convolution operation of GCN on the neighbor nodes with dissimilar features.

Although DGMP has achieved good performance in pan-cancer driver gene prediction, it can be improved from the following two aspects. First, DGMP averages the SNVs, CNVs, DNA methylation and gene expression of all samples to obtain 4-moics features to represent the genes for each cancer, which will ignore the specific characteristics from individual cancer patients. Second, DGMP just simply concatenated three embedding feature vectors from DGCN and one embedding feature vector from MLP, while the contributions of these four feature vectors may be different. Thus, we can consider using the weighted fusion way to improve the performance of DGMP.

## CRediT author statement

**Shao-Wu Zhang**: Conceptualization, Methodology, Investigation, Formal analysis, Writing - review & editing. **Jing-Yu Xu**: Conceptualization, Methodology, Software, Investigation, Writing - original draft. **Tong Zhang**: Investigation, Analysis, Writing – partial original draft. All authors read and approved the final manuscript.

## Acknowledgments

We would like to thank Drs. Yan Li and Wei-Feng Guo for their work on the manuscript revision. This work was supported in part by the National Natural Science Foundation of China (Grant Nos. 62173271, 61873202 to SWZ)

