## Supplementary material for "DGMP: Identifying Cancer Driver Genes by Jointing DGCN and MLP from Multi-Omics Genomic Data": Table S5

**Table S5 AUPR and AUC of DGMP, EMOGI, NRFD, HotNet2, MutSigCV, PageRank and DeepWalk+SVM on DawnNet and STRING PPI network in 5CV test.**

| Method | DawnNet**^@^** | | STRING PPI**^@@^** | |
| --- | --- | --- | --- | --- |
|  | AUPR | AUC | AUPR | AUC |
| DGMP | **0.761** | **0.886** | **0.964** | **0.971** |
| EMOGI | 0.617 | 0.825 | 0.914 | 0.947 |
| NRFD | 0.641 | 0.801 | 0.693 | 0.736 |
| PageRank | 0.595 | 0.724 | 0.666 | 0.712 |
| DeepWalk+SVM | 0.589 | 0.799 | 0.685 | 0.783 |
| HotNet2 | 0.621 | 0.726 | 0.673 | 0.712 |
| MutSigCv | 0.315 | 0.593 | 0.465 | 0.563 |

^@^ 693 cancer driver genes and 1763 non-cancer driver genes are used to train prediction models.

^@@^ 734 cancer driver genes and 1152 non-cancer driver genes are used to train prediction model
